# Fasten with Pipes

**DOI:** 10.1101/2023.10.30.564744

**Authors:** Lee S. Katz, Henk C. Den Bakker

## Abstract

Fasten is a versatile bioinformatics tool suite developed to address gaps in the manipulation of sequence data files in the fastq format. Existing tools often lack support for interleaved fastq files and Unix-style pipe file control. Fasten provides a solution by allowing bioinformaticians to efficiently work with paired-end sequence data, facilitating data transformation, and offering competitive speeds. This suite, implemented in Rust, offers a comprehensive manual, continuous integration, and seamless integration into the command line with Unix pipes. Fasten is poised to become an invaluable resource for routine command-line bioinformatics tasks, enhancing the accessibility and flexibility of data processing in the field.

## Background

There are still many gaps in basic command line bioinformatics for standard file formats. Bioinformaticians have been able to use many tools to manipulate sequence data files in the fastq format, such as seqkit (Shen 2016), seqtk (Li 2023) or FASTX-Toolkit (Gordon 2014). These tools only accept paired end (PE) sequence data when split into multiple files per sample. Additionally, these tools do not always allow for Unix-style pipe file control. Sometimes they require explicity input/output options instead of using stdin and stdout. However, some bioinformaticians prefer to combine PE data from a single sample into one file using the interleaved fastq file format, but this format is not always well supported in mainstream tools. Here, we provide Fasten to the community to address these needs.

## Materials

We leveraged the Cargo packaging system in Rust to create a basic framework for interleaved fastq file manipulation. Each executable reads from stdin and prints reads to stdout and only performs one function at a time. The core executables perform these fundamental functions: 1) converting to and from interleaved format, 2) converting to and from other sequence file formats, 3) ‘straightening’ fastq files to a more standard 4-line-per-entry format.

There are 20 executables including but not limited to read metric generation, read cleaning, kmer counting, read validation, and regular expressions for interleaved fastq files.

We have also taken advantage of Rust to make comprehensive and standardized documentation. Continuous integration was implemented in GitHub Actions for unit testing, containerizing, and benchmarking. Benchmarking was performed against other mainstream packages using hyperfine using 100 replicates and 2 burn-ins (Peter 2023).

## Results

Documentation, the container, and code are available at GitHub. Benchmarking results were graphed into Figure 1.

**Figure 1.**
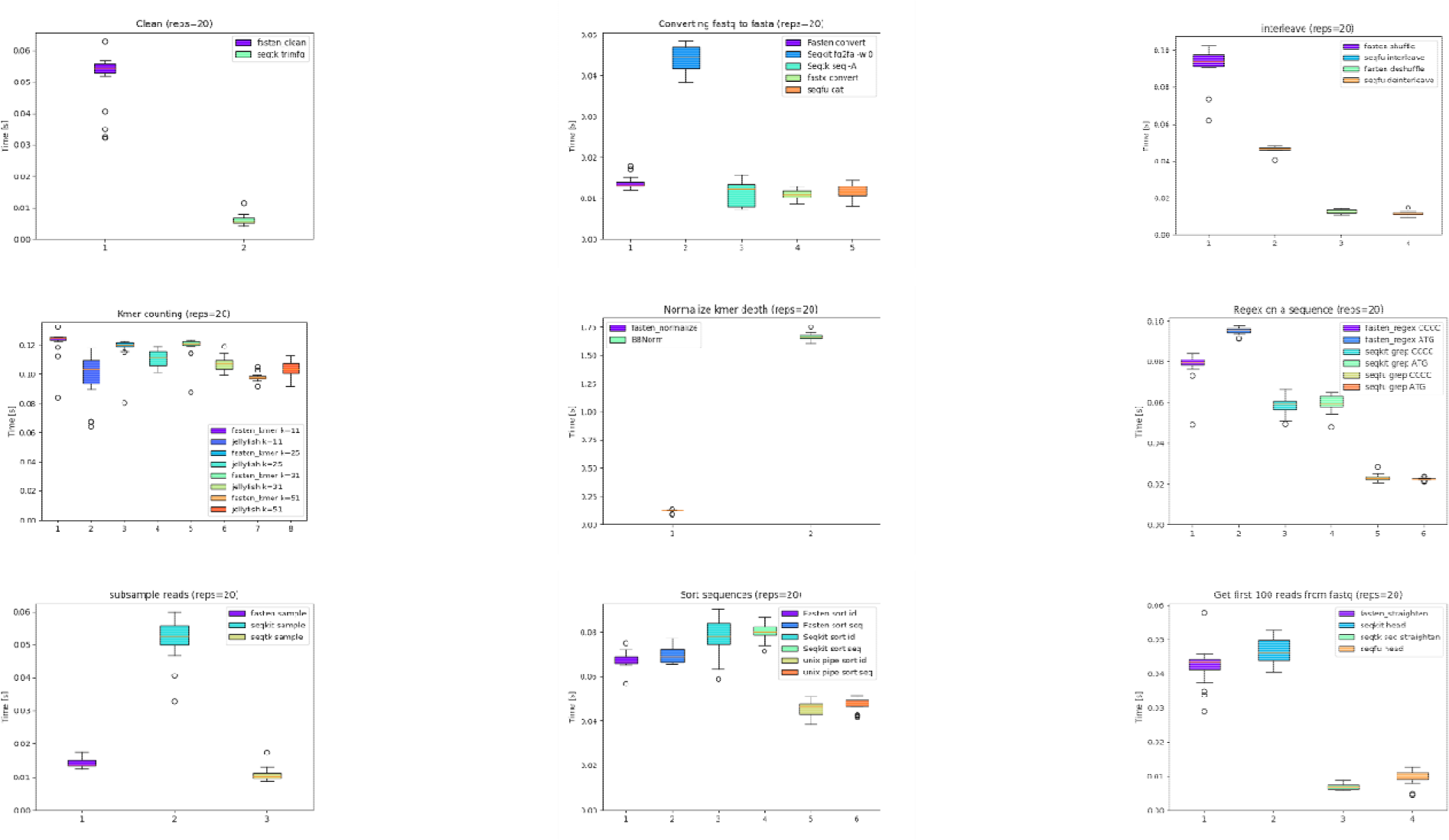
Benchmarks comparing fasten with other analagous tools. From left to right, then to bottom: Trimming with a minimum quality score; converting fastq to fasta; interleaving R1 and R2 reads; kmer counting; normalizing read depth using kmer coverage; Searching for a sequence in a fastq file; downsampling reads; sorting fastq entries by either sequence or ID; and converting nonstandard fastq files to a format whose entries are four lines each, and selecting the first 100.

## Conclusions

Fasten is a powerful manipulation suite for interleaved fastq files, written in Rust. We benchmarked Fasten on several categories. It has strengths as shown in Figure 1 but it does not occupy the fastest position in all cases. Its major strengths include its competetive speeds, Unix-style pipes, paired-end handling, and the advantages afforded by the Rust language including documentation and stability.

Fasten touts a comprehensive manual, continuous integration, and integration into the command line with unix pipes. It is well poised to be a crucial module for daily work on the command line.

## Acknowledgements

Thank you, John Phan, for creating the Docker container.

